## Supplemental Figures for "Choroid-resident macrophages maintain local vasculature and RPE integrity and spontaneously regenerate following depletion"

**This PDF file includes:**

Figures S1 to S6

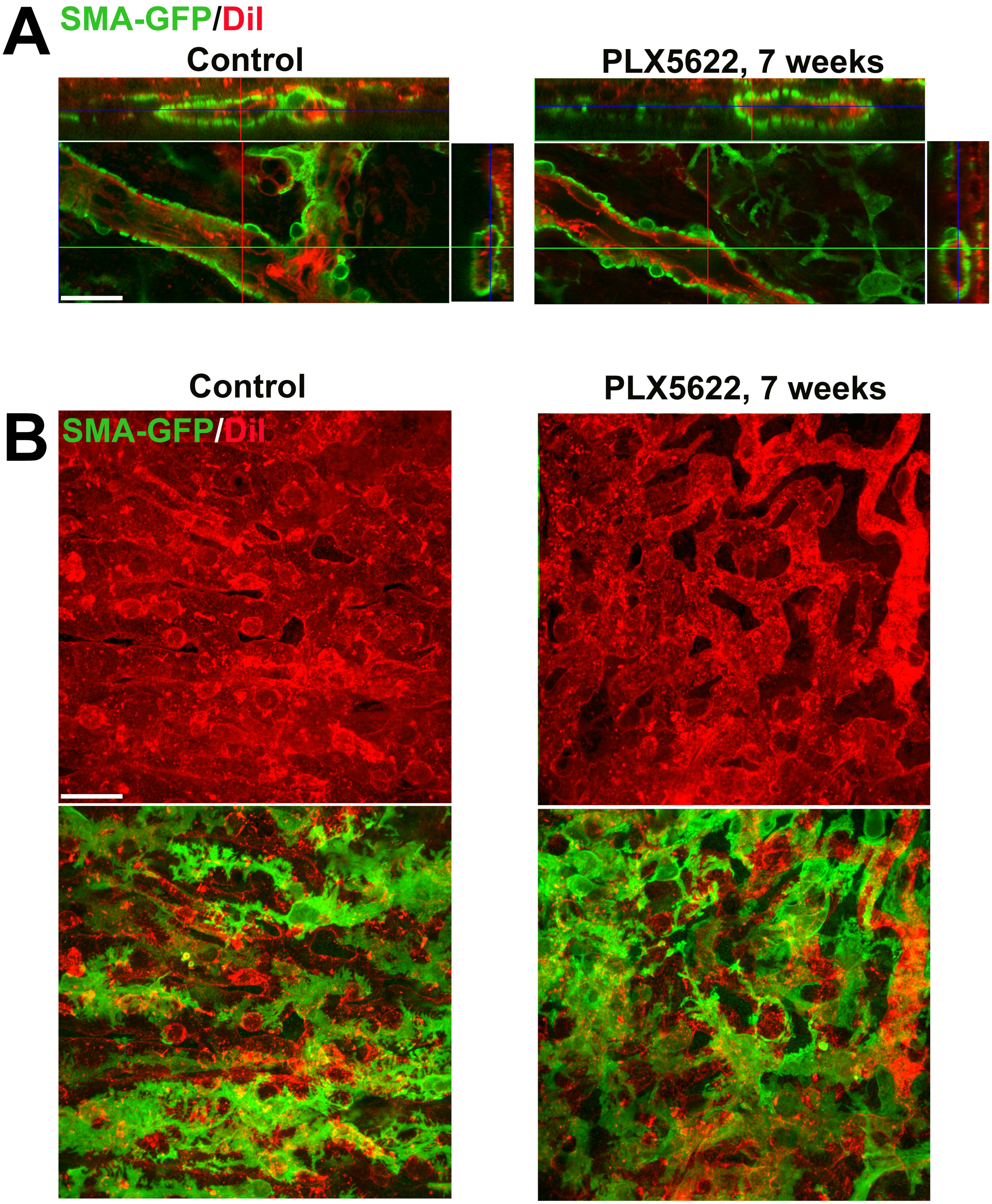


**Fig. S1. Long-term depletion of resident choroidal macrophages does not result in marked depletion or disorganization of smooth muscle actin (SMA)-expressing perivascular smooth muscle cells or choriocapillaris pericytes.** Albino transgenic mice possessing a transgene containing an α-SMA promoter driving the expression of green fluorescent protein (GFP) were used to visualize perivascular cells of the choroidal vasculature. The choroidal vasculature was visualized by systemic DiI perfusion (red). **(A)** Perivascular smooth muscle cells of adult 3-month old mice administered diet containing PLX5622 for 7 weeks to deplete choroidal macrophages were compared with those in controls (age-matched mice fed standard diet). Orthogonal views of confocal images demonstrated complete coverage of large choroidal vessels (red) with contiguous perivascular cells (green) that were largely unaltered with choroidal macrophage depletion. **(B)** Confocal images of the choriocapillaris visualized enface demonstrated a similar presence and coverage of ramified pericytes on the sclerad surface of choriocapillaris vasculature in depleted and control animals. Scale bars = 20 µm.

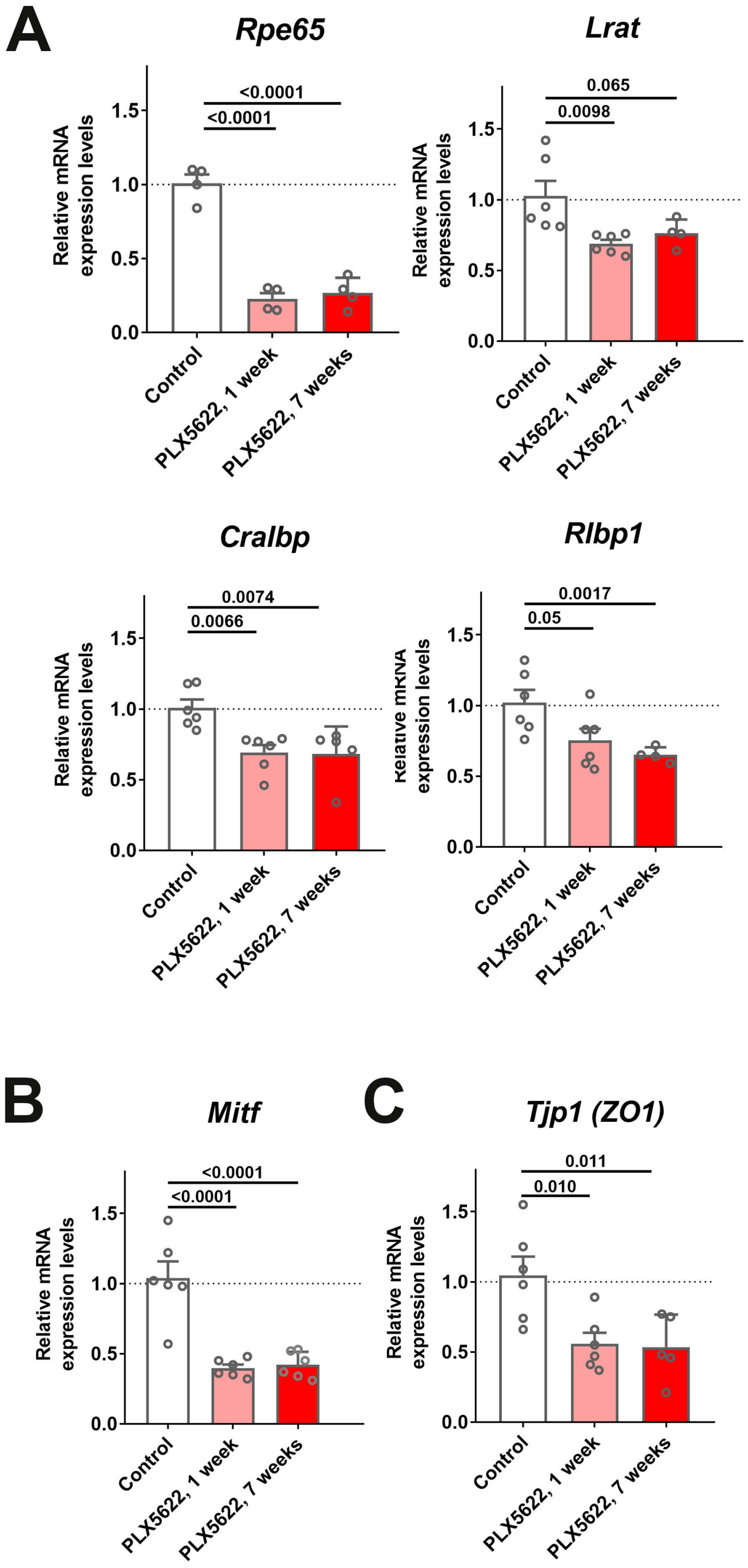


**Fig. S2. mRNA expression levels of genes related to RPE function are altered following transient and sustained depletion of choroidal macrophages.** Changes in mRNA expression of RPE-specific genes in sclerochoroidal tissue following 1 and 7 weeks of continuous PLX5622 administration were evaluated with RT-PCR. Age-matched untreated animals served as controls. **(A)** RPE genes related to visual cycle function demonstrated general decreases; *Rpe65* expression was markedly decreased beginning at 1 week of treatment, with smaller relative changes were observed for *Lrat*, *Cralbp*, and *Rlbp1*. *Mitf,* a transcriptional factor important for RPE differentiation **(B)**, and *Tjp1* (ZO1), a tight-junctional protein between RPE cells **(C)**, also showed significant decreases. P values correspond to comparisons relative to control made using a 1-way ANOVA test, n = 4-6 animals each treatment group.

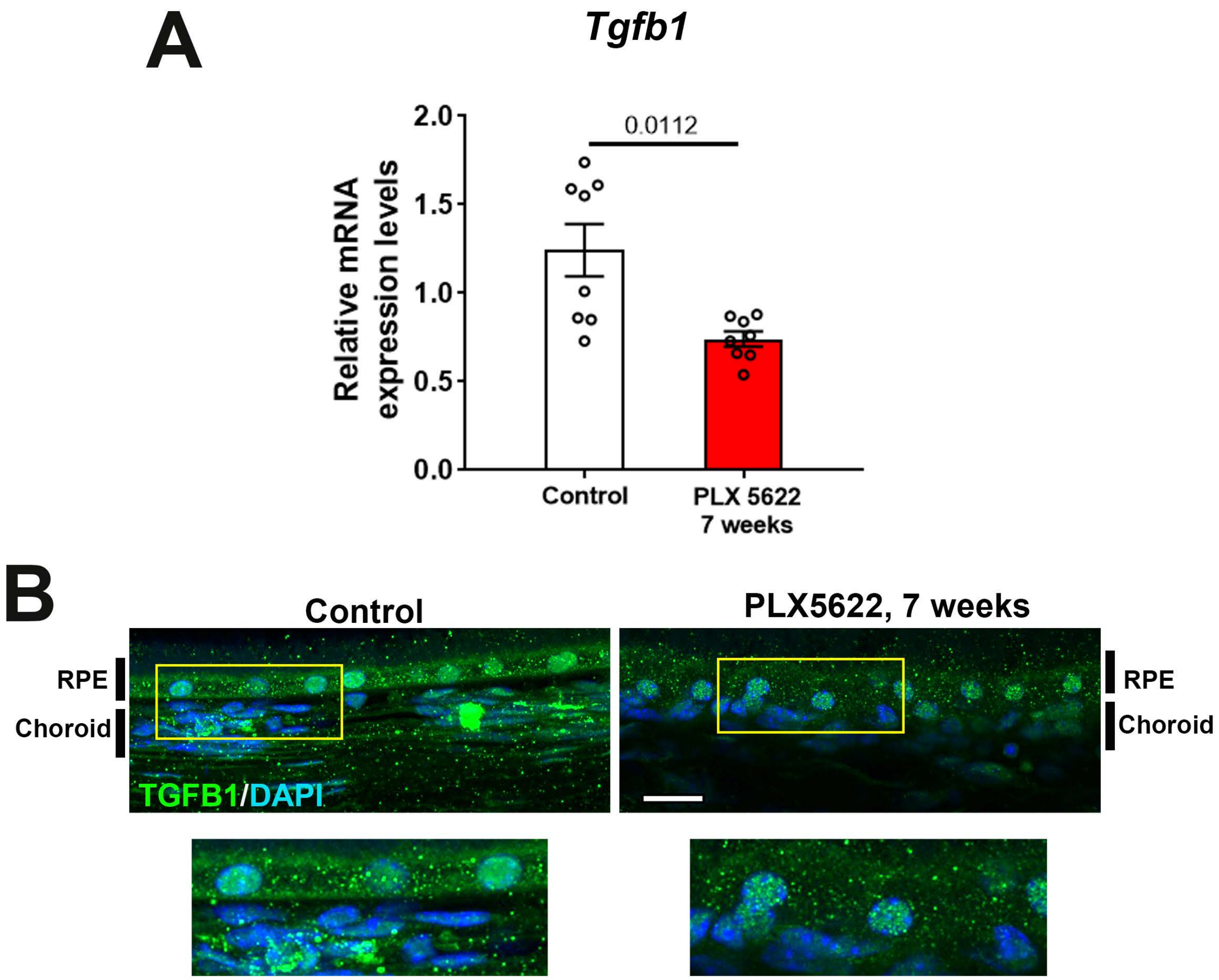


**Fig. S3. mRNA expression levels of Tgfb1 were altered following sustained depletion of choroidal macrophages.** Changes in mRNA expression of Tgfb1 in sclerochoroidal tissue were evaluated with RT-PCR following 7 weeks of continuous PLX5622 administration. Age-matched untreated animals served as controls. mRNA level of *Tgfb1* **(A)** was significantly reduced following 7 weeks of macrophage depletion. Immunohistochemical localization demonstrated decreased TGFβ1 immunopositivity in the RPE layer and choroid following macrophage depletion **(B).** P values represent comparison to control level using an unpaired t-test with Welch's correction. Scale bars = 20 μm.

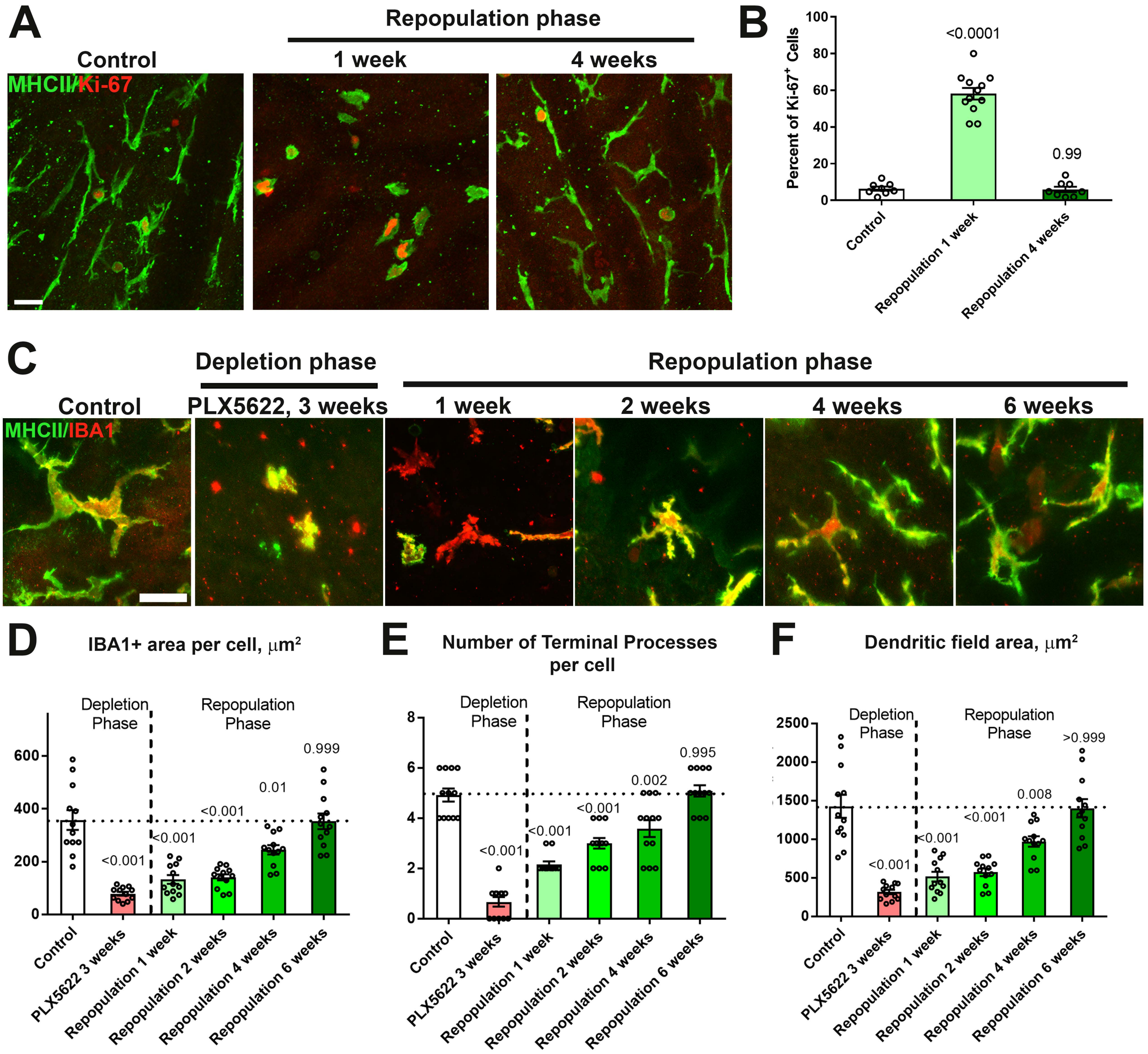


**Fig. S4. Repopulating choroidal macrophages demonstrate proliferation in situ and progressive morphological maturation. (A, B)** Ki67 immunopositivity was absent in MHCII+ repopulating macrophages in undepleted control animals but was prominent at 1 week of repopulation. The proportion of Ki67+ in MHCII+ cells declined to control levels by 4 weeks. Scale bar = 20μm. **(C)**High magnification images of choroidal macrophages showing deramified morphologies during depletion and a progressive recovery of ramified morphologies during macrophage repopulation. Scale bar = 20μm. Quantitative morphological analyses of repopulating macrophages showed progressive enlargement of cell size (defined as the area of IBA1-immunopositiviey per cell) **(D),** increasing ramification (in terms of number of terminal processes per cell) **(E)**, and increasing dendritic field size (defined as the area of the bounding polygon around each cell) **(F)**. Full recapitulation of these morphological measures back to baseline levels was achieved at 6 weeks into the repopulation phase. P values represent comparisons to control levels, 1-way ANOVA with Tukey's correction for multiple comparisons.

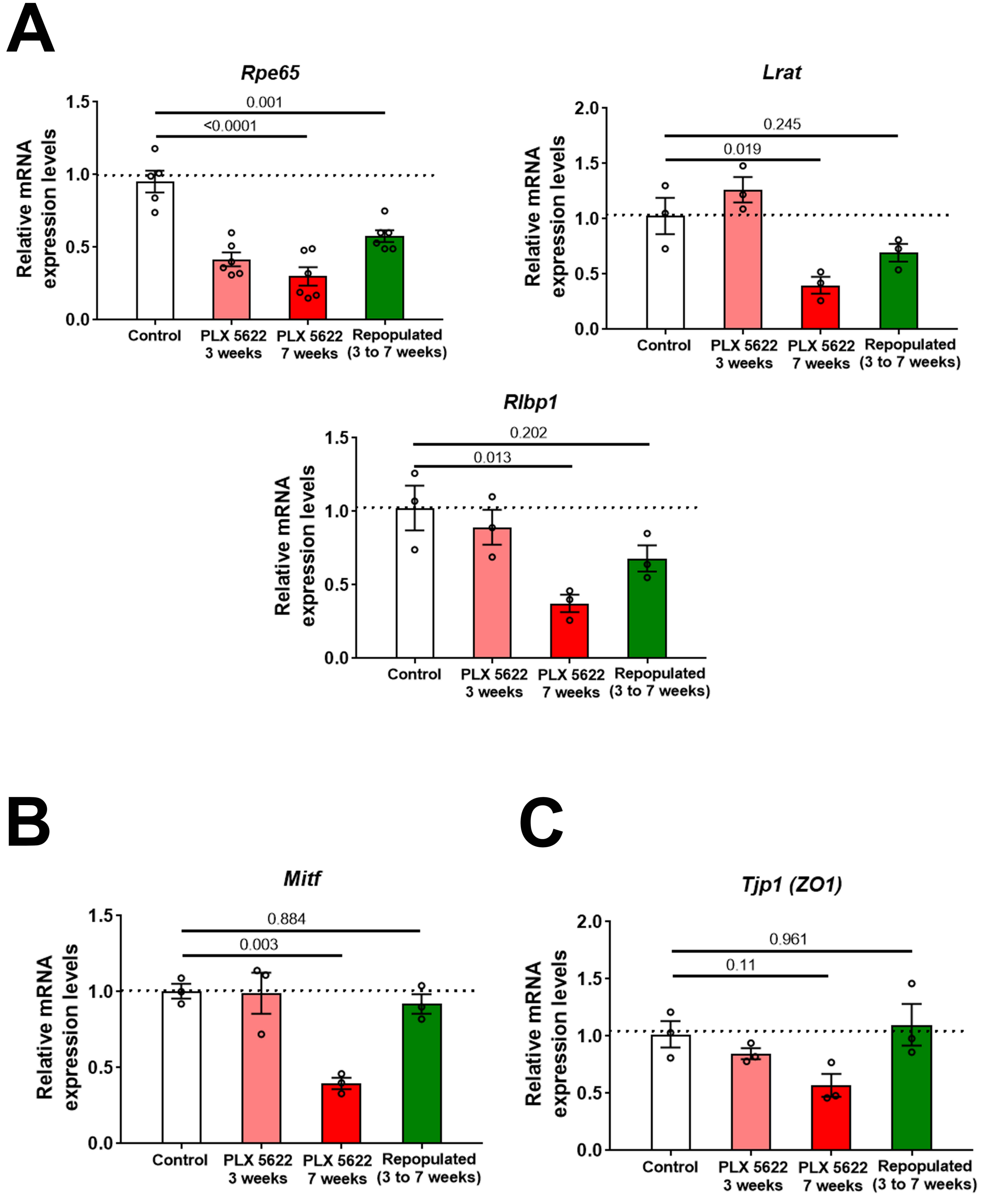


**Fig. S5. Macrophage depletion-associated changes in RPE genes expression are ameliorated with macrophage repopulation.** Changes in mRNA expression of RPE-specific genes in sclerochoroidal tissue from the following experimental groups were evaluated with RT-PCR: (1) untreated age-matched controls, (2) macrophage-depleted for 3 weeks, (3) macrophage-depleted for 7 weeks (continuous depletion group), and (4) macrophage-depleted for 3 weeks, followed by 4 weeks of macrophage repopulation (depletion-repopulation group). **(A)** RPE genes related to visual cycle function (*Rpe65, Lrat, Rlpb1*), **(B)** *Mitf,* and **(C)** *Tjp1/Zo1*, demonstrated decreases with depletion that were ameliorated or reversed with macrophage repopulation. P values correspond to comparisons relative to control made using a 1-way ANOVA test, n = 3 animals each treatment group.

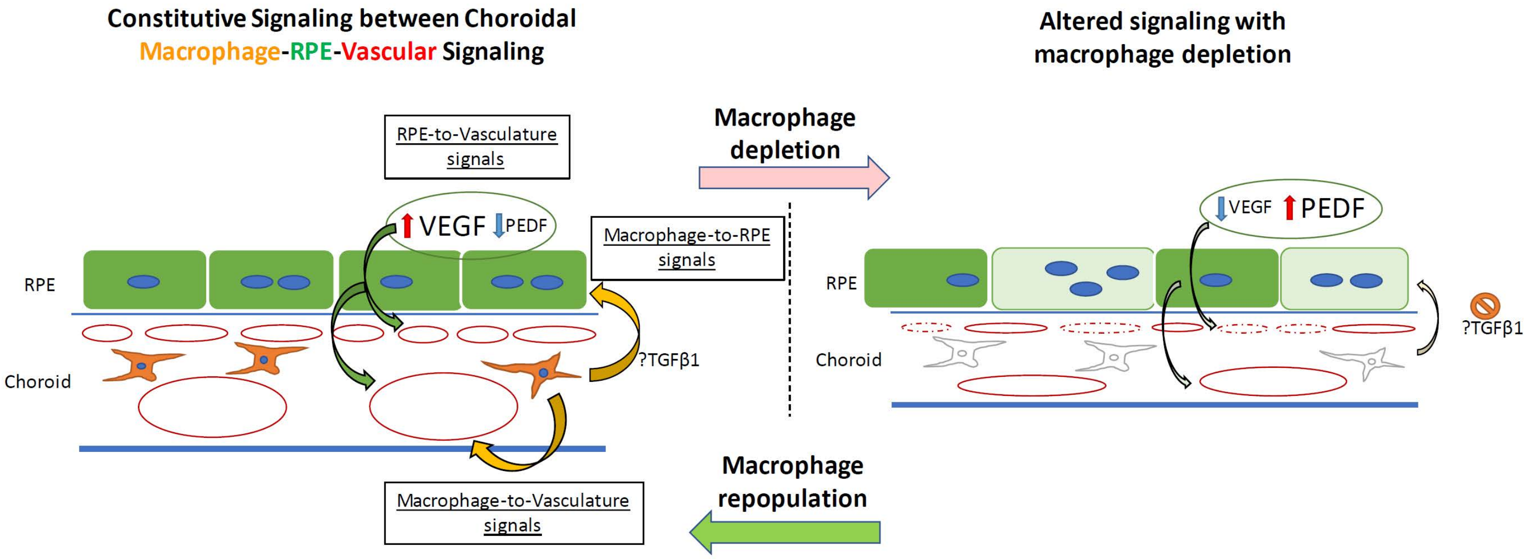


**Fig. S6. Schematic showing putative signaling between choroidal macrophages, vasculature, and the RPE layer in the outer retina under normal conditions and with choroidal macrophage depletion. (Left)** In the normal adult choroid, angiogenic signals (VEGF, PEDF) secreted from the RPE cell layer regulate local choroidal vascular structure. Choroidal macrophages produce potential trophic signals that influence RPE structure and angiogenic secretion (e.g. TGFB1), secondarily influencing vascular maintenance. Potential uncharacterized angiogenic signals may also originate from choroidal macrophages to impinge onto choroidal vasculature to maintain them. **(Right)** With the ablation of choroidal macrophages, macrophage trophic signals are diminished, leading to RPE cell structure changes and altered balanced of angiogenic factor production by RPE cells. These induce choroidal vascular atrophy, reducing choroidal thickness and decreasing choriocapillaris coverage.
